# An expanded gene catalog of the mouse gut metagenome

**DOI:** 10.1101/2020.09.16.299339

**Authors:** Jiahui Zhu, Huahui Ren, Huanzi Zhong, Xiaoping Li, Yuanqiang Zou, Mo Han, Minli Li, Lise Madsen, Karsten Kristiansen, Liang Xiao

## Abstract

High-quality and comprehensive reference gene catalogs are essential for metagenomic research. The rather low diversity of samples used to construct existing catalogs of mouse gut metagenomes limits the size and numbers of identified genes in existing catalogs. We therefore established an expanded gene catalog of genes in the mouse gut metagenomes (EMGC) containing >5.8 million genes by integrating 88 newly sequenced samples, 86 mouse-gut-related bacterial genomes and 3 existing gene catalogs. EMGC increases the number on non-redundant genes by more than one million genes compared to the so far most extensive catalog. More than 50% of the genes in EMGC were taxonomically assigned and 30% were functionally annotated. 902 Metagenomic species (MGS) assigned to 122 taxa are identified based on the EMGC. The EMGC-based analysis of samples from groups of mice originating from different animal providers, housing laboratories and genetic strains substantiated that diet is a major contributor to differences in composition and functional potential of the gut microbiota irrespective of differences in environment and genetic background. We envisage that EMGC will serve as an efficient and resource-saving reference dataset for future metagenomic studies in mice.

## Introduction

Mouse models are among the most widely used animal models for biomedical studies to decipher the complex interplay between the gut microbiota and host phenotypes (Nguyen et al., 2015; Justice & Dhillon, 2016; Perlman, 2016; Hugenholtz & de Vos, 2018). Amplicon sequencing of the 16S rRNA gene has been widely used for analyses of the gut microbiota due to low costs and short analysis cycle. However, the taxonomic information is in most cases limited to the genus level, and amplicon sequencing generally provided limited information on function (Morgan & Huttenhower, 2014; Wang & Jia, 2016b). Key to the use of mouse model for detailed functional analyses of the gut microbiota is the availability of comprehensive catalogs of microbial genes and derived metagenomic species (MGSs)/metagenome-assembled genomes (MAGs). The first catalog of genes in the mouse gut microbiome included 2.6 million nonredundant genes from fecal samples of 184 mice (Xiao et al., 2015). Subsequent studies further explored the diversity and functional potential of the mouse gut microbiota by isolating and sequencing an increasing number of bacterial strains from the mouse gut (Wannemuehler et al., 2014; Lagkouvardos et al., 2016; Brugiroux et al., 2017; Liu et al., 2020;) and establishing a Mouse Intestinal Bacterial Collection (miBC), depositing bacterial strains and associated genomes from the mouse gut (Lagkouvardos et al., 2016). Recently, Lesker *et al*. generate an integrated mouse gut metagenome catalog (iMGMC), comprising 4.6 million unique genes and 830 high-quality MAGs, and by linking MAGs to reconstructed 16S rRNA gene sequences they provided a pipeline enabling improved prediction of functional potentials based on 16S rRNA gene amplicon sequencing (Lesker et al., 2020).

Here we constructed an expanded mouse gut metagenome catalog (EMGC) by integrating 3 published gene catalogs, including the gene catalog of mouse gut metagenome (MGGC) released in 2015 comprising 2,571,074 genes (Xiao et al., 2015), a Feed & Diet Gene Catalog for mice (FDGC)(Xiao et al., 2017), the integrated mouse gut metagenome catalog (iMGMC)(Lesker et al., 2020), 72 available sequenced mouse gut-related bacterial genomes (Wannemuehler et al. 2014; Lagkouvardos et al., 2016; Brugiroux et al., 2017; Liu et al., 2020), 14 high-quality genomes assembled from published sequencing data of isolates (Lagkouvardos et al., 2016), and 88 newly shotgun sequenced samples. Our new non-redundant reference gene catalog comprises 5,862,027 genes and was annotated by NR (released on 5^th^ Jan 2019) and KEGG (release 89) databases (Kanehisa & Goto, 2000). Finally, we generated 902 MGSs from the gene abundance profiles for 326 laboratory mice of EMGC, and compared these MGSs with the high-quality MAG collection (Lesker et al., 2020) and the recent collection of bacteria isolated from the mouse gut (Liu et al., 2020). By combining these individual datasets we increased the number of sequenced bacterial genes of the mouse gut microbiome by more than one million genes, and significantly increased the mapping ratio of read obtained by shotgun sequencing of samples from mouse gut fecal samples providing a resource for future studies on the mouse gut microbiota.

## MATERIALS and METHODS

### Data acquisition

DNA from 88 stool samples of C57BJ/6J wild-type male mice, collected from the laboratory of BGI-Wuhan, were extracted and shotgun sequenced using the BGISEQ-500 platform and PE100 sequencing as describe (Fang et al., 2018). An optimized sequencing quality filter for the cPAS-based BGISEQ platform, OAs1 (Fang et al., 2018), was applied in the quality control step, followed by host removal with SOAP2 (Li et al., 2009). Individual assembly of metagenomic reads was done using metaSPAdes v3.14.0 (parameters: -k 49, other parameters were set to be default)(Kultima et al., 2012; Nurk et al., 2017). Genes were predicted by MetaGeneMark (W. Zhu et al., 2010) from metagenome-assembled contigs with length >500bps and filtered by length >100bps. Redundant predicted genes were removed by CD-HIT (v4.5.7, parameters: -G 0 -n 8 -aS 0.9 -c 0.95 -d 0 -r 1 -g 1) (Fu et al., 2012) in order to generate a sub mouse gut gene catalog (PMGC).

Raw reads of 43 unassembled bacterial genomes from the miBC (EBI Project ID PRJEB10572) were downloaded from EBI and filtered by Trimmomatic (v 0.39)(Bolger et al., 2014). Draft genomes were assembled separately by SPAdes (-k 29,39,49,69 –careful) (Bankevich et al., 2012; Nurk et al., 2013) and filtered by the CheckM (Parks et al., 2015). After assessment using the criteria Completeness >90% and Contamination <5%, the remaining 14 genomes were used for gene prediction by GeneMarkS-2 (Lomsadze et al., 2018).

A total of 72 genomes, including 24 sequenced strains of miBC(Lagkouvardos et al., 2016), 8 genomes of the Altered Schaedler Flora(Wannemuehler et al., 2014) (PRJNA175999-176003, 213740, 213743) and 40 genomes of mGMB (Liu et al., 2020) (released before 26^th^ Feb 2019, PRJNA486904), as well as their coding sequences (CDSs) and translated CDSs were all downloaded from the NCBI RefSeq database and the Integrated Microbial Genomes (IMG) database(Chen et al., 2019). We gathered CDSs from 86 bacterial genomes and filtered out genes smaller than 100bps. We clustered CDSs using CD-HIT (v4.5.7, parameters: -G 0 -n 8 -aS 0.9 -c 0.95 -d 0 -r 1 -g 1) (Fu et al., 2012) establishing a gene catalog termed Mouse Intestinal Cultured Bacteria gene set (MiCB). Detailed information on the included genomes is provided in Supplementary Table 3.

The five public mouse-related microbial datasets used in this study include: (i) 184 host-free sequenced mice gut microbiomes (EBI Project ID PRJEB7759) and the catalog of mouse gut metagenome (MGGC, GigaDB http://dx.doi.org/10.5524/100114); (ii) 54 sequenced mice gut microbiomes (EBI Project ID PRJEB10308) and the related gene catalog (FDGC, GigaDB http://dx.doi.org/10.5524/100271); (iii) 830 high-quality de-replicated MAGs and the iMGMC catalog from the iMGMC study (GitHub repository https://github.com/tillrobin/iMGMC); (iv) 40 sequenced mouse fecal metagenomes from EBI Project PRJEB6996; (v) 34 sequenced mice cecum metagenomes from MG-RAST Project mgp6153.

### Construction of EMGC and selection of new genes

All downloaded gene were filtered by length >100bps and integrated to construct the EMGC using CD-HIT (Fu et al., 2012) (parameters: -G 0 -n 8 -aS 0.9 -c 0.95 -d 0 -r 1 -g 1). The output of EMGC clusters was analyzed to generate a list for new genes in EMGC that are not present in iMGMC and MGGC. Metagenomes were mapped to gene catalogs by SOAP2 (parameters: -m 0 -x 1000 -c 0.95). Mapping rates between groups and catalogs were compared by Wilcoxon rank-sum test. The profile of relative gene abundances for the 326 laboratory mice (see Supplementary Table 1 for an overview of these mice) was calculated based on the method of Qin et al.(Qin et al., 2012) using EMGC. Richness estimation by the Chao2 index and incidence-based coverage estimator (ICE) was calculated based on the gene abundance profiles (Li et al., 2014).The occurrence and average abundance of new genes were calculated using the relative gene abundance profiles.

### Taxonomic and functional annotation

Genes predicted from metagenome assemblies were taxonomically annotated by Kaiju (Menzel et al., 2016) using the NCBI-NR database (released on Jan 5^th^, 2019) and the parameters of the program were set to ‘-a greedy -e 5 -E 0.01 -v -z 4’. For genes from mouse gut-related bacterial genomes, we kept the original taxonomic information of the genomes and assigned them to the corresponding genes. All genes were searched against KEGG (Kanehisa & Goto, 2000) (version 89) by BLAST (Altschul et al., 1990; Gish & States, 1993) for functional annotation. Parameters were set to ‘-e 0.01 - F T -b 100 -K 1’ and results were filtered by score >60 (20). BLAST results were turned into functional annotation based on the information provided by KEGG to create a gene-KO list for the generation of KO relative abundance profiles. The taxonomic and functional information of new genes of EMGC were extracted for further analysis.

### Evaluation of the effect of diet on the gut microbiota

The differences between sample groups regarding mapping rate were assessed by Wilcox rank-sum test (R ggpubr package). The significance for mapping rate was set at p <0.05. To evaluate the consistent effect of diet among providers and mouse strains on taxonomic and functional composition of gut metagenomes, samples from 7 groups fed high-fat (HF) or low-fat (LF) diets and representing different providers and mouse strains were selected (Supplementary Table 1). The calculation of relative abundance profiles for taxa and KO was based on Qin et al. (2012). Permutational multivariate analysis of variance (PERMANOVA) test was applied to determine the influence of diet on gut metagenomes within different groups. The Shannon index of the relative abundance profiles was used to estimate alpha diversity of the samples. Principal Coordinates Analysis (PCoA) of selected samples was performed based on the relative abundance profiles using Bray-Curtis distance (R ape4 package) to visualize the effect of diet on the bacterial composition of the gut microbiota. Wilcox rank-sum test was used for analysis of differences of genera and KO relative abundance profiles. P-value adjustment was applied for multiple hypothesis testing using the Benjamin-Hochberg (BH) method. A BH-adjusted P-value < 0.05 was considered as statistical significance.

### Metagenomic species clustering

The gene relative abundance profiles of 326 laboratory mice were clustered using the co-abundance Canopy algorithm (Nielsen et al., 2014). Co-abundance genomes (CAGs) that were present in >90% samples were chosen and CAGs with >700 genes were considered as Metagenomic Species (MGSs)(Nielsen et al., 2014). CAGs and MGSs were assigned to a given taxon when >50% genes belonged to that specific taxon (Nielsen et al., 2014). The taxonomic distribution of MGSs was calculated using the R package ‘phytool’(Revell, 2012). The Shannon Index was calculated based on the MGS profiles of samples from the 7 selected mice groups and PCoA was generated based on the same profile.

All MGSs were searched against the 830 high quality Metagenome-Assembly Genomes (MAGs) and the 115 mGMB genomes by MUMmer3 (v3.23) (Kurtz et al., 2004) for calculation of MUMi values (Bäckhed et al., 2015; Deloger et al., 2009). If the MUMi value for two items was >0.54, then these two items were recognized as the same species (Bäckhed et al., 2015). The result of the comparison between MGSs and MAGs was hierarchical clustered by R package ‘hclust’ with weighted pair group method with averaging (WPGMA) and then visualized in a cladogram with annotation by a R package ‘ggtree’(Yu, 2020; Yu et al., 2017). The result of the comparison between MGSs and mGMB genomes is presented in Supplementary Table 9.

## Results

### Construction and evaluation of EMGC

Fecal samples from 88 C57BL/6J male mice were sequenced using the BGISEQ-500 platform providing 1,098 gigabases (Gb) high-quality host-free data with an average of 12.47 Gb per sample (Supplementary Table 2) and a catalog comprising 2,602,584 non-redundant genes (PMGC). We next used 72 mouse gut related bacterial genomes (Wannemuehler et al., 2014; Lagkouvardos et al., 2016; Brugiroux et al., 2017; Liu et al., 2020;) from IMG and NCBI Refseq and 14 high-quality genomes (completeness >90% and contamination <5%) assembled from reads accessible from PRJEB10572 (Lagkouvardos et al., 2016) (Supplementary Table 3) to generate a Mouse Gut Cultured Bacteria gene set (MiCB). Together with MGGC (Xiao et al., 2015), FDGC (Xiao et al., 2017) and iMGMC (Lesker et al., 2020), downloaded from GigaDB and the GitHub repository, respectively, all gene catalogs were integrated to construct an expanded non-redundant mouse gut bacterial gene catalog (EMGC)(Figure 1). The expanded catalog comprises 5,862,027 genes, which is more than twice the number of genes in the MGGC (Xiao et al., 2015) and one million genes more than the iMGMC catalog (Lesker et al., 2020) (Table 1). Thus, 18.93 % of the genes in EMGC are not represented in either the iMGMC or MGGC (Supplementary Figure 1).

**Table 1.**
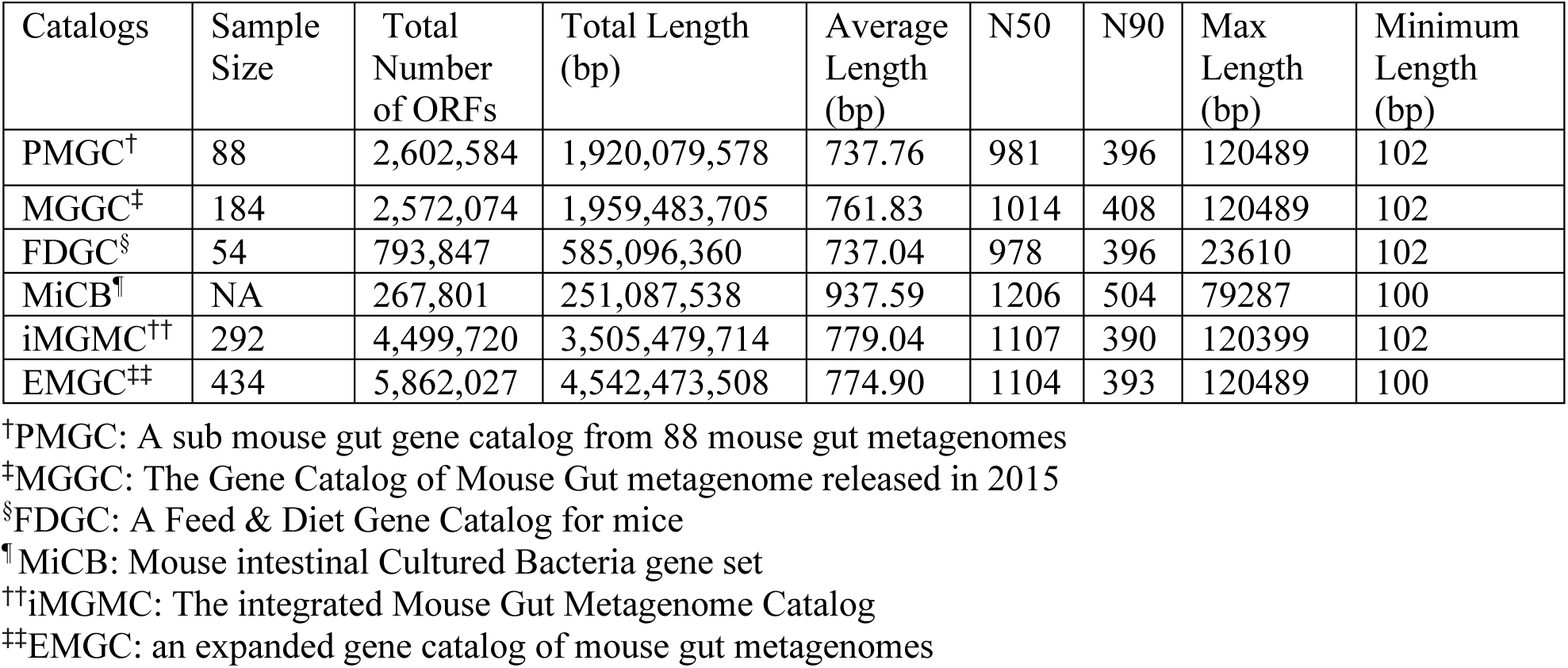
General Features of gene catalogs

**Figure 1.**
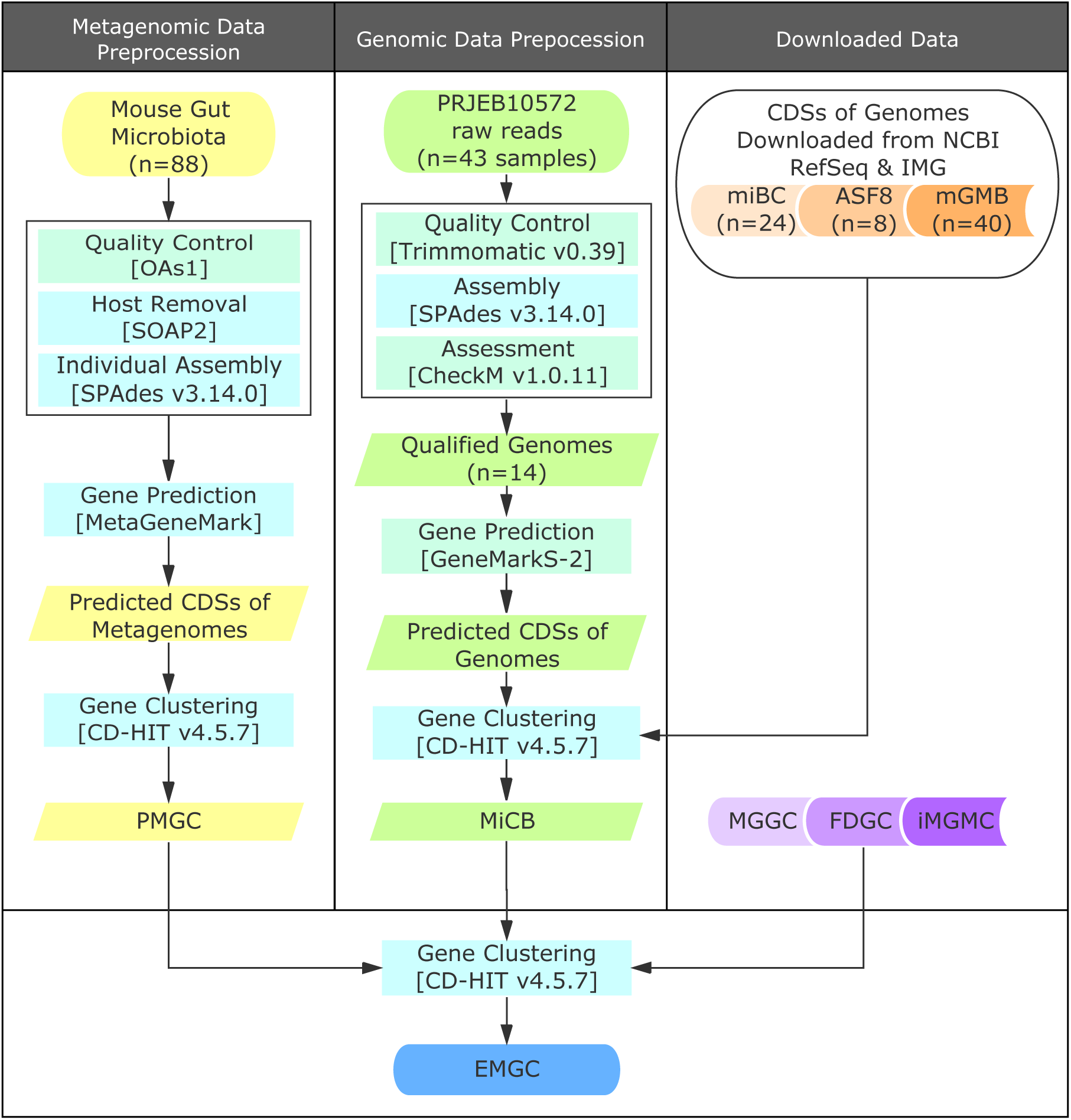
Construction of the EMGC. Metagenomic sequencing data of 88 mouse gut metagenomes were processed by the pipeline as displayed to generate non-redundant genes for PMGC. Unassembled strains of miBC (under BioProject PRJEB10572) were assembled and filtered by genome quality (completeness >90%, contamination <5%) of assembled genomes. Qualified genomes were used for gene prediction. CDSs from assembled genomes and downloaded genomes were gathered and clustered to MiCB. PMGC and MiCB alongside with 3 downloaded genesets, FDGC, MGGC and iMGMC, and were merged to generate EMGC.

To compare the performance of EMGC relative to MGGC and iMGMC, we mapped sequencing reads from the FDGC, MGGC, and PMGC studies to the three catalogs. Of the sequencing reads from PMGC, 55.72% were mapped to MGGC and 56.56 % to the iMGMC, whereas the EMGC allowed mapping of 79.52% of the reads (Figure 2A), close to the maximum achievable mapping rate in prokaryotes (Li et al., 2014).

**Figure 2.**
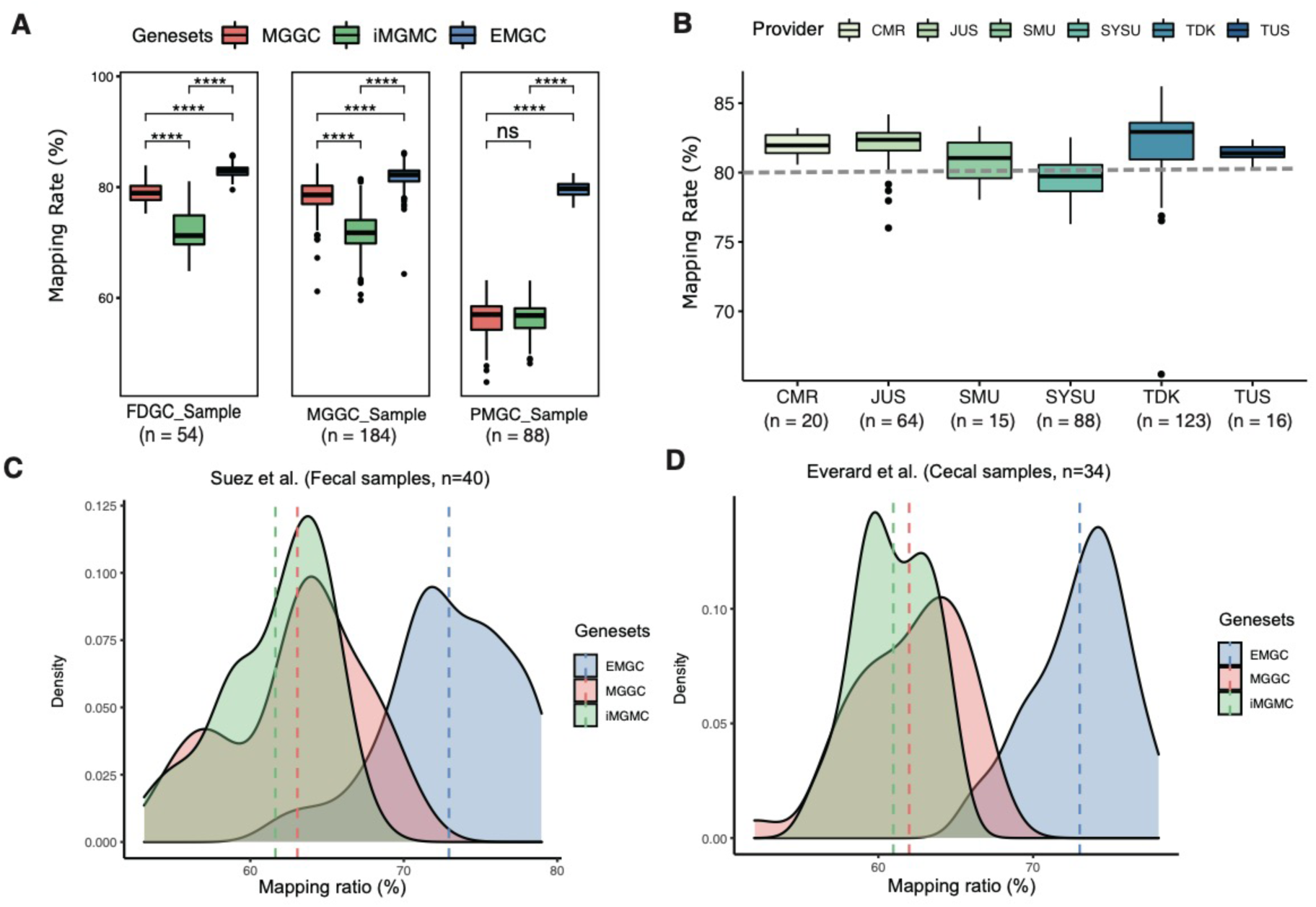
Performance of the EMGC. (A) Comparison of mapping rates between MGGC, iMGMC and EMGC. **** represents p values <0.0001 by Wilcox rank-sum test. (B) Display of mapping rates among samples’ providers, including the Wallenberg Laboratory for Cardiovascular and Metabolic Research (CMR), the Jackson Laboratory in US (JUS), Laboratory Animal Center of Southern Medical University (SMU), Laboratory Animal Center of Sun Yat-Sen University (SYSU), Taconic in Denmark (TDK) and US (TUS). Dashed line represents a mapping rate of 80%. (C) & (D) Density curve for the mapping rate of fecal metagenomes and cecal metagenomes which were not included in gene catalog construction. Dashed lines represented the average values of the mapping rate of each gene catalog.

Comparing mapping rates of reads from the 326 fecal samples obtained from different mouse strains and providers to the EMGC demonstrated that mice from the Laboratory Animal Center at Sun Yat-Sen University exhibited a lower mapping rate than samples from the other providers where the median mapping rates were higher than 80% (Figure 2B). Median mapping rates of reads obtained from all mice strains were also higher than 80% (Supplementary Figure 2A). Richness estimated by Chao2 indicated that our EMGC covered 98.24% of the genes in the 326 fecal samples (Supplementary Figure 2B) whereas ICE suggested that 97.49% of the genes were covered.

To further evaluate the quality of the EMGC, we mapped the metagenomic data obtained from 40 fecal samples from control mice and mice that had consumed non-caloric artificial sweeteners (Suez et al., 2014) and metagenomic data obtained from 34 cecal samples from control mice and mice treated with prebiotic (Everard et al., 2014). For the reads obtained in the study of Suez et al., 63.07% mapped to the MGGC and, 61.04 % to iMGMC, whereas 72.94% mapped to the EMGC (Figure 2C; Supplementary Table 4). For the read obtained from the study of Everard et al., 61.99% mapped to MGGC and 60.97% mapped to iMGMC, but 73.01% of the reads mapped to the EMGC (Figure 2D; Supplementary Table 5). Together these results demonstrate a significantly increase mapping rate of reads using the EMGC as a reference.

### Taxonomic and functional characteristics of EMGC

We taxonomically annotated the genes of EMCG using Kaiju (Menzel et al., 2016) and the NCBI NR database to provide an overview of the taxonomical composition visualized by a Krona plot (Ondov et al., 2011). This plot revealed that 67% of the genes could be annotated (Supplementary Figure 3). We assigned 54.20% of the genes to the phylum level and 44.30% of the genes to the family level (Supplementary Figure 4A, 4B). We next annotated the genes in the EMGC to the KEGG (release 89) database (Kanehisa & Goto, 2000) and identified 6571 KEGG functional orthologs (KO) and 292 KEGG pathways (Supplementary Figure 5).

To further examine the quality of the EMGC, we calculated the occurrence frequency and average abundance of the 1,109,381 genes not present in the previous MGGC and iMGMC. As shown in Figure 3A, 40.41% of these genes exhibited an occurrence frequency and mean abundance higher than 0.1 and 10^−8^, respectively. We also extracted taxonomic and functional information of these new genes. Annotation of the genes not present in the MGGC at the species level revealed that the top 5 species could be assigned to *Oscillibacter sp. 1-3, Firmicutes bacterium ASF500, Acetatifactor muris, bacterium 1xD42-67* and *Eubacterium plexicaudatum* (Figure 3B), all isolated from the mouse gut based on information from NCBI BioSample Database. In relation to functions, the general distribution of KEGG pathways in these additional genes is similar to the overall distribution in EMGC (Figure 3C). We identified 190 KOs in EMGC which are not present in either MGGC or iMGMC. Furthermore, 46 KEGG pathways are covered by additional KOs (Supplementary Table 6) in EMGC. For 4 KEGG pathways, including betalain biosynthesis, xylene degradation, neomycin, kanamycin and gentamicin biosynthesis and lipopolysaccharide biosynthesis, we found that more than 5% of the additional KOs are only represented in EMGC compared to iMGMC (Figure 3D, Supplementary Table 6).

**Figure 3.**
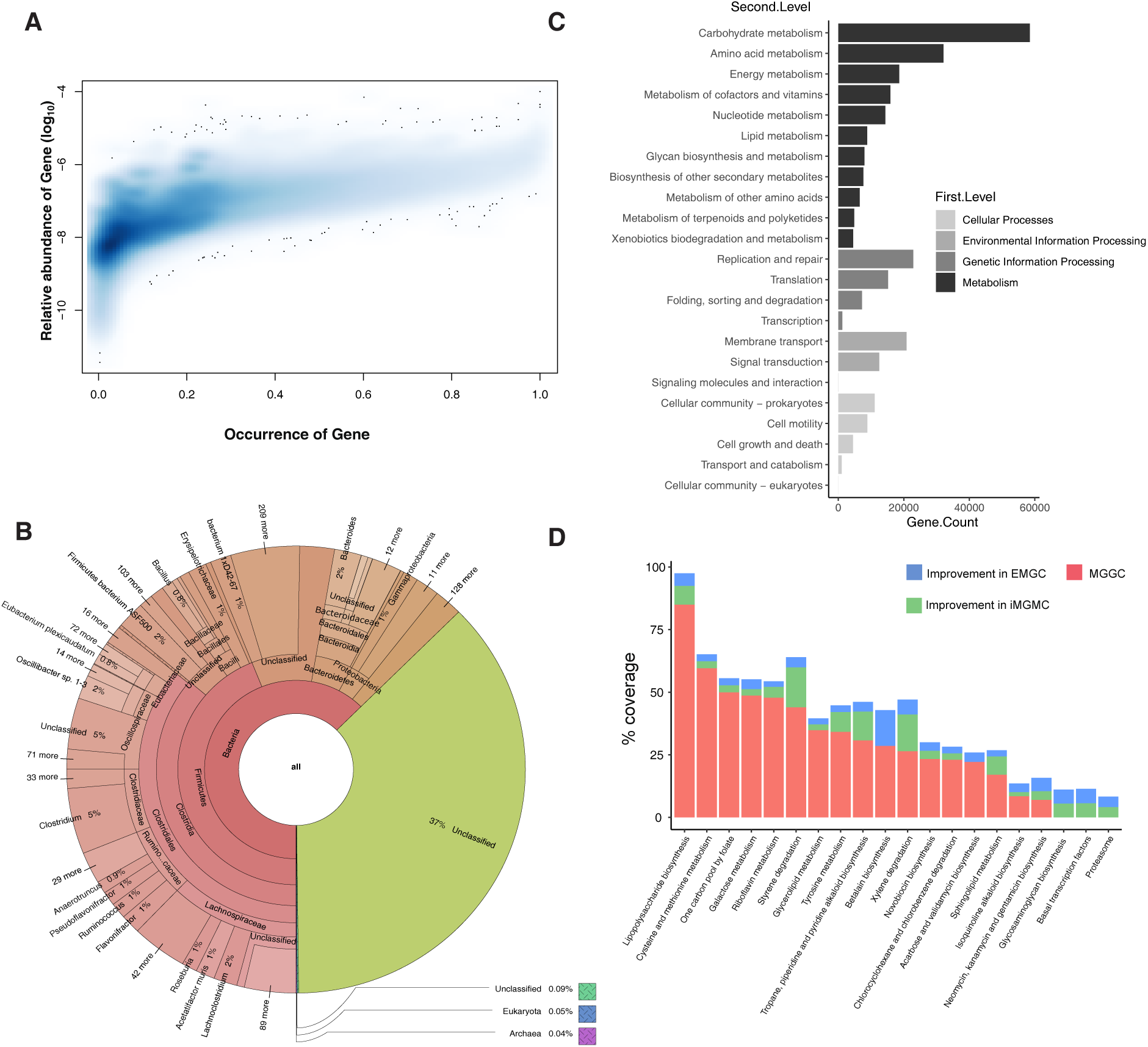
Description of new genes included in EMGC. (A) 2D density histogram showing the distribution of occurrence and mean relative abundance of new genes. (B) A general display of taxonomic composition of new genes by Krona. (C) Frequency of functional pathways associated with the new genes. (D) Stacked histogram of KO coverage of functional pathways improved in EMGC, compared to MGGC and iMGMC. Coverage is calculated as (*Annoatated ko numbers/Total ko numbers*) × 100% in a given pathways.

### Changes in the microbiota composition and functional potential

We have earlier reported that the mouse gut metagenome is affected by animal providers and housing as well as strain and diet (Xiao et al., 2015). In order to investigate to what extent diet affected the gut metagenomes independently of strain and providers, we selected 7 groups (G1-G7) of samples from different strains from different providers fed a low-fat (LF) diet or a high-fat (HF) diet and housed in the same facility (Details in Supplementary Table 1). We estimated the impact of diet on the variation of gut microbiota based on the relative abundance profiles of genera and KOs using PERMANOVA. The analyses indicated that diet explained at least 33.9% (P value=0.003) of the total variation at the genus level and 47.3 % (P value=0.006) at the KO level (Figure 4A, Supplementary Figure 6A). Compared to mice fed a LF diet, mice fed a HF diet exhibited an increase in alpha diversity at the genus level independent of housing laboratories, strains and providers (Figure 4B). In contrast, at the KO level alpha diversity in mice fed a LF diet generally, except for group 7, exhibited an increased diversity compared to mice fed a HF diet (Supplementary Figure 6B). Principal coordinate analysis (PCoA) similarly confirmed that the diet strongly influenced the genus profile (Figure 4C), and the KO profile (Supplementary Figure 6C).

**Figure 4.**
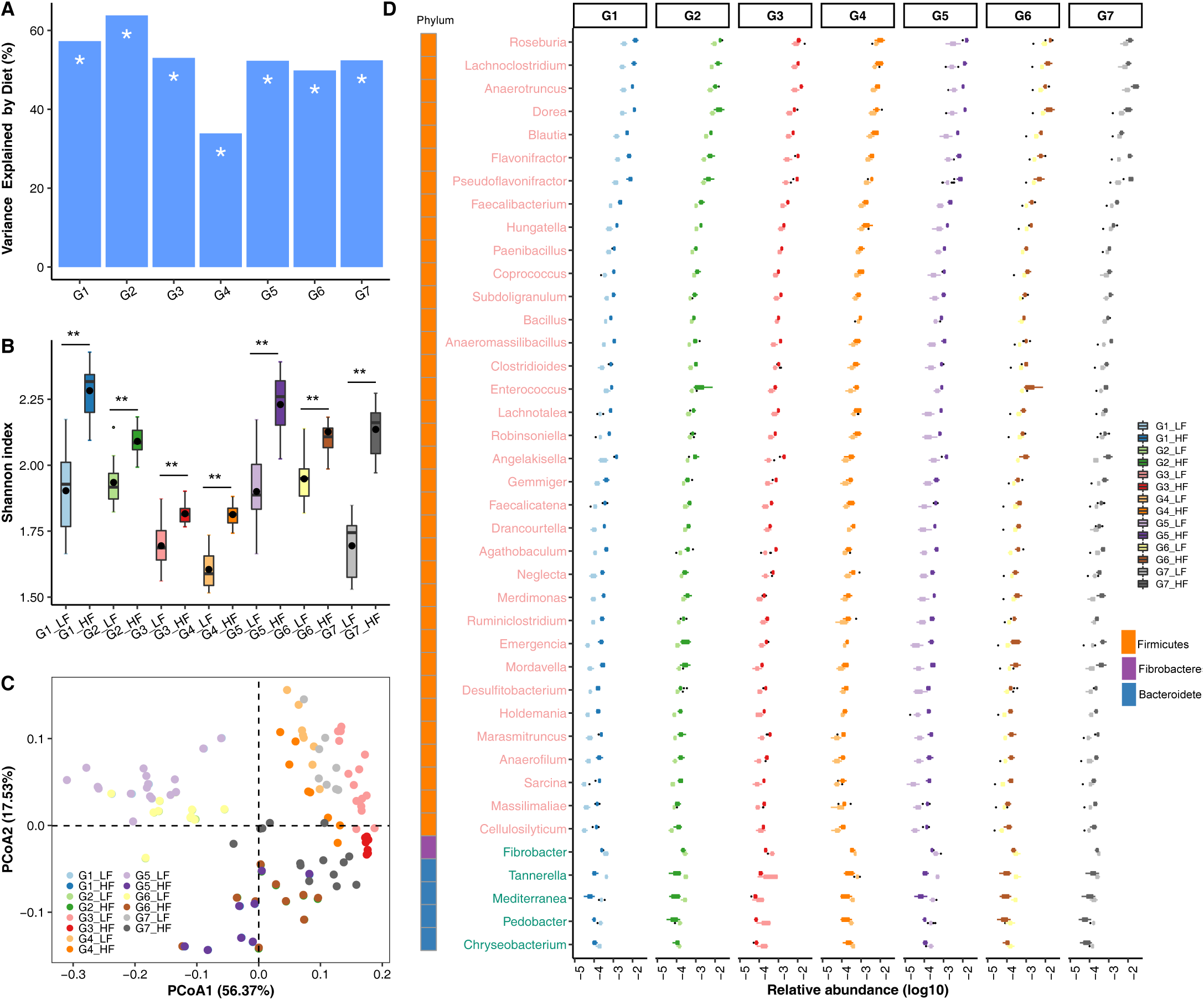
The influence of diet on the composition of the microbiota. (A) PERMANOVA analysis to estimate the influence of diet on the composition of gut metagenomes among all 7 sample groups. G1: C57/BL6 mice provided by Taconic Denmark (TDK) and hosted in National Institute of Nutrition and Seafood Research of Norway (NIFES); G2: Sv129 mice provided by TDK and hosted in NIFES; G3: C57/BL6 mice provided by the Jackson Laboratory in US (JUS) and hosted by Pfizer-I; G4: C57/BL6 mice provided by Taconic US (TUS) and hosted in Pfizer-I; G5: C57/BL6 mice provided by TDK and hosted in the University of Copenhagen (KU); G6: Sv129 mice provided by TDK and hosted in KU; G7: C57/BL6 mice provided by the Laboratory Animal Center of Sun Yat-Sen University (SYSU). (B) Shannon index (** represents p value <0.01, Wilcox rank-sum test) and (C) PCoA based on genus profile for 7 groups fed the HF and LF diet. (D) Genera differently enriched (BH-adjusted p-value <0.05, Wilcox rank-sum test; relative abundance >1e-5) in mice fed the HF and LF diet among all 7 groups. Genera in light red represent genera enriched in HF fed mice, while genera in light green represent genera enriched in LF fed mice.

To further examine diet-induced changes, we examined genera and KOs enriched in samples from either HF or LF fed mice by Wilcoxon rank-sum test. As shown in Figure 4D, in all 7 groups the 36 genera found at higher abundance in samples from HF fed mice belong to Firmicutes, whereas four genera within the Bacteroidetes phylum and one genus within the Fibrobacteres phylum were found at higher abundance in samples from LF fed mice (Figure 4D, Supplementary Table 7). However, whereas 324 KOs were enriched in LF-fed mice, only 233 KOs were detected at higher abundance in HF than LF fed mice (Supplementary Table 8). To investigate which taxa contributed to the disparate response to LF and HF diet at the taxonomy and the functional levels, we identified the taxa at the phylum level that contributed to the enrichment of KOs. Whereas genera within the Bacteroidetes phylum were the main contributor to KOs enriched in LF-fed mice, genera within the Firmicutes phylum contributed in HF-enriched KOs (Supplementary Figure 7).

### Construction of metagenomic species

We identified 902 Metagenomic Species (MGSs, >700 genes) using the relative gene abundances based on 326 fecal samples obtained from different mouse strains and providers using MGS Canopy Clustering and taxonomic annotation as described previously (Nielsen et al., 2014). The 902 MGSs were assigned to 122 taxa (Supplementary Figure 8; Supplementary Table 9). We also generated MGS profiles for the 7 groups of mice fed a LF or a HF diet. The Shannon indices and PCoA plot revealed a clear effect of diet, independent of mouse strain and provider (Supplementary Figure 9A & 9B).

We next compared the 902 MGSs with the 830 high-quality non-redundant MAGs generated in the iMGMC project (Lesker et al., 2020) and 115 bacterial genomes from the mGMB project (Liu et al., 2020). 559 (61.97%) MGSs were classified as the same species as the MAGs from the iMGMC project (Maximal unique match index (MUMi) value (Bäckhed et al., 2015; Deloger et al., 2009; J. Li et al., 2020) >0.54, Supplementary Table 9). As shown in Figure 5, Firmicutes and Bacteroides were the most prevalent phyla among all MGSs and MAGs. We also identified 56 MGSs representing genomes of species from the mGMB project, and of these, 8 MGSs could be identified as mGMB genomes, but not as MAGs (Supplementary Table 9). Of note, for more than one third of the MGSs we were unable to identify corresponding entities in the MAG collection or in the cultured genomes collection.

**Figure 5.**
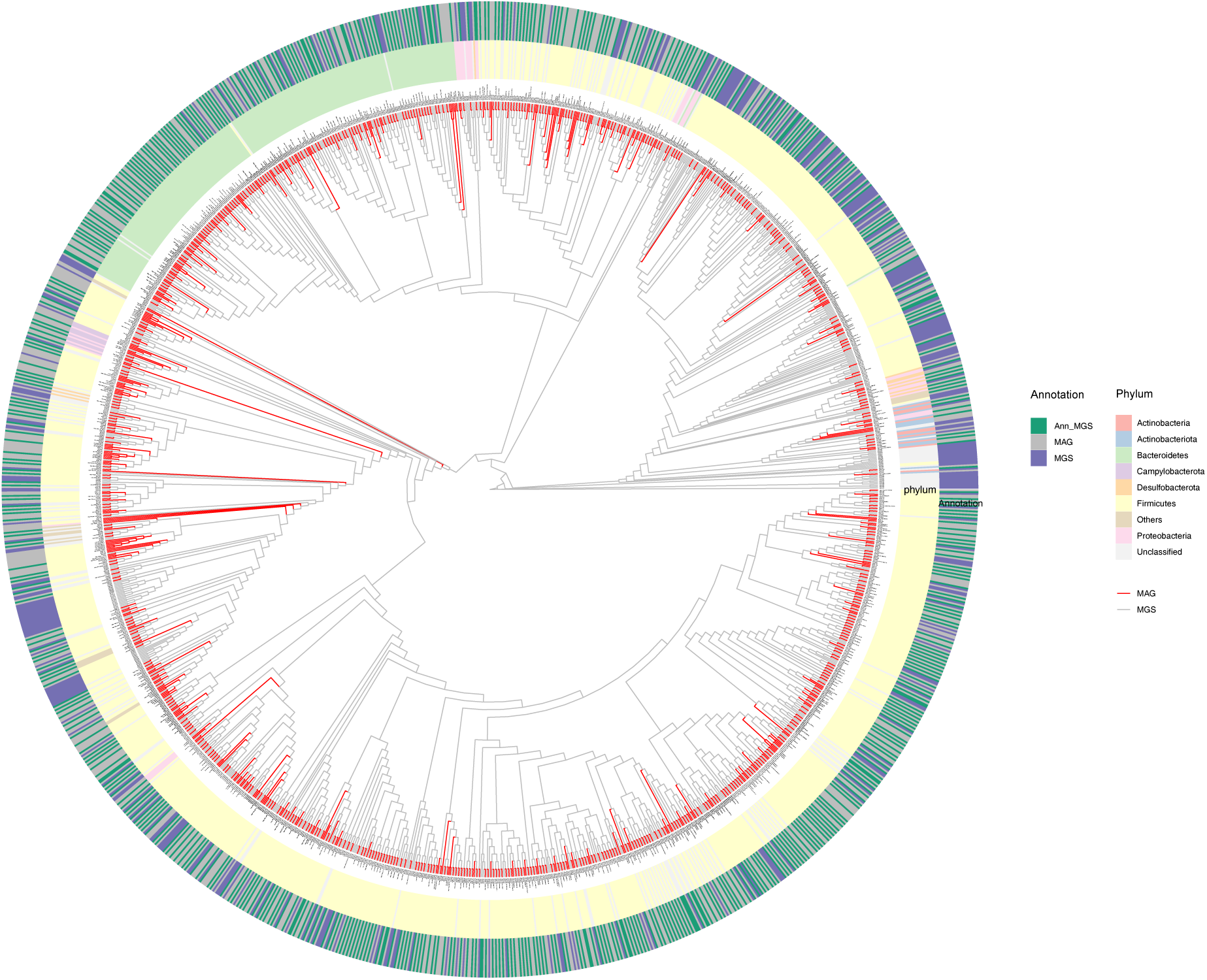
Phylogenetic tree of the 902 MGSs and 830 high-quality iMGMC MAGs. MUMi distance for MGSs and MAGs were used to construct the phylogenetic tree using hierarchical clustering. MAGs are shown as red branches and MGSs as grey branches. The outer ring shows the relation between MGSs and MAGs. MGSs which have a MUMi value >0.54, are marked as “Ann_MGS” in green blocks, otherwise in purple blocks. MAGs are all in grey blocks. Colored blocks in the inner cycle indicate phyla assigned to MGSs and MAGs.

## Discussion

The EMGC represents the most comprehensive catalog of genes in the mouse gut microbiome. It covers samples from feces and cecum, different mouse strains fed different diets, obtained from different providers and housed in different laboratories. The majority of the genes identified in this study were assigned to known species, which might improve the coverage of known species and the detection of low abundant taxa. The improvement in KO coverages of a number of pathways will enhance the functional characterization of the mouse gut microbiota. In addition, the analysis of samples from different mouse strains from different animal providers and different housing laboratories confirms the pronounced effect of diets on the taxonomic and functional composition of the gut microbiota.

In spite of the increased number of genes in EMGC, there are still some limitations. The sample size and variation of sample types in the EMGC are still small. The majority of samples included in EMGC were collected from C57/BL6 mice, which might affect the applicability in studies on other laboratory mouse strains and wild-caught mice. Many additional confounding factors than those addressed in the present study will most probably impinge on the gut microbiota (Hugenholtz & de Vos, 2018; Laukens et al., 2016; Nguyen et al., 2015; Perlman, 2016). It therefore seems important to include more samples from other mouse strains to gain further insights into the effect of confounding factors, which may lead to pronounced variability in the mouse gut microbiota, which again might limit the reproducibility of biomedical research using mouse models (Laukens et al., 2016; Stappenbeck & Virgin, 2016). Although both culture-independent and culture-dependent studies on the mouse gut microbiota have been carried out to improve the understanding of host-microbe interaction in mouse models, the majority of mouse gut metagenome members still remains relatively uncharacterized (Lagkouvardos et al., 2016; Lesker et al., 2020; Liu et al., 2020; Xiao et al., 2015).

Even though more and more metagenomic analysis methods are being developed at an increasing speed, gene catalogs are still necessary resources, not only to provide reliable and consistent taxonomic annotation, but also to reduce the gap between phylogenetic and functional biases (Li et al., 2014; Nayfach & Pollard, 2016). The high-quality reference gene catalog, EMGC, together with the 902 metagenomic species, is able to support and improve accurate metagenome-wide association analyses using mouse models which may assist in functional characterization of observed correlation between the microbiota composition and functional potential in relation host phenotypes.

## Acknowledgements

This work is funded by the National Natural Science Foundation of China (Grant No. 81670606), the National Key Research and Development Program of China (No. 2018YFC1313800) and the Shenzhen Engineering Laboratory of Detection and Intervention of Human Intestinal Microbiome (No. DRC-SZ [2015]162). This work was supported by China National GeneBank and CNSA (CNGB Nucleotide Sequence Archive) of CNGBdb. We thank the sequencing and bioinformatic staff of BGI-Shenzhen, especially Dr. Dan Wang, Ms. Ying Xu, Ms. Fangming Yang, Mr. Jie Zhu, Dr. Jinghong Yu and Dr. Guangwen Luo for help and advice in our work.

## Data availability

The host-free data that was sequenced in this study have been deposited into CNSA (CNGB Nucleotide Sequence Archive) of CNGBdb with accession number CNP0000619 (metagenomic sequencing reads and assemblies of 88 mice samples with the dataset of EMGC).

## Author contributions

J. Z., X.L. and Liang X. managed the project. X.L., Y.Z., H.Z., M.L. and Liang X. supervised the project. X.L. provided samples. J.Z., H.R., H.Z., G.L. and Liang X. designed the analysis. J.Z., H.R. and G.L. performed the analysis. J.Z. wrote the paper. J.Z., H.R., H. Z., X.L., Y. Z, M. H., M. L., K.K. and Liang X. revised the paper.

## Competing financial interests

The authors declare no competing financial interests.

## Notes

### Competing Interest Statement

The authors have declared no competing interest.

