## Supplementary Figures for "An expanded gene catalog of the mouse gut metagenome"

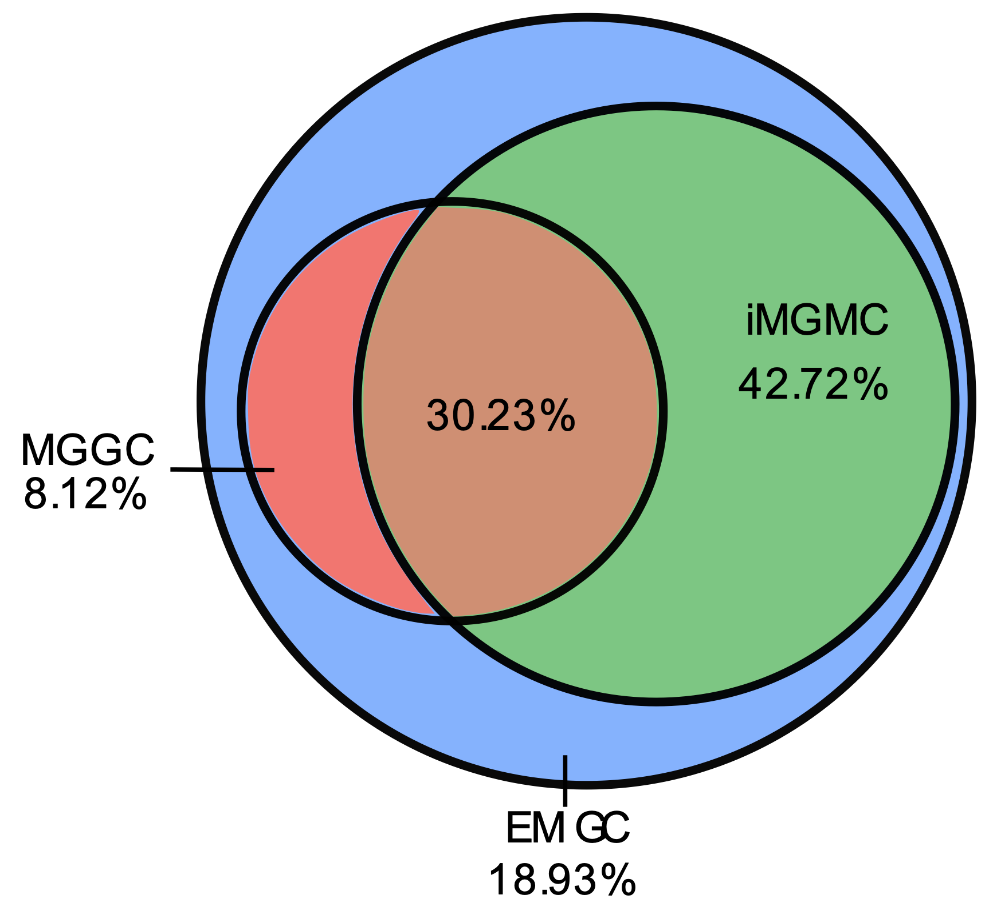

**Supplementary Figure 1** Unique and shared Genes comparing MGGC, iMGMC and EMGC.

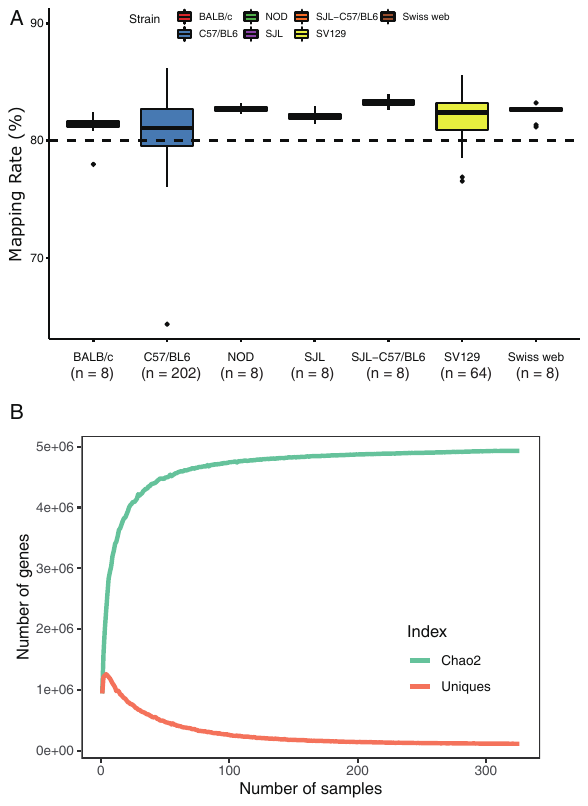

**Supplementary Figure 2** (A) Mapping Rate based on the EMGC grouped by Mouse Strains. Grid line indicates a mapping rate of 80%. (B) Rarefaction Curve of Chao2 Index and Unique Gene Count.

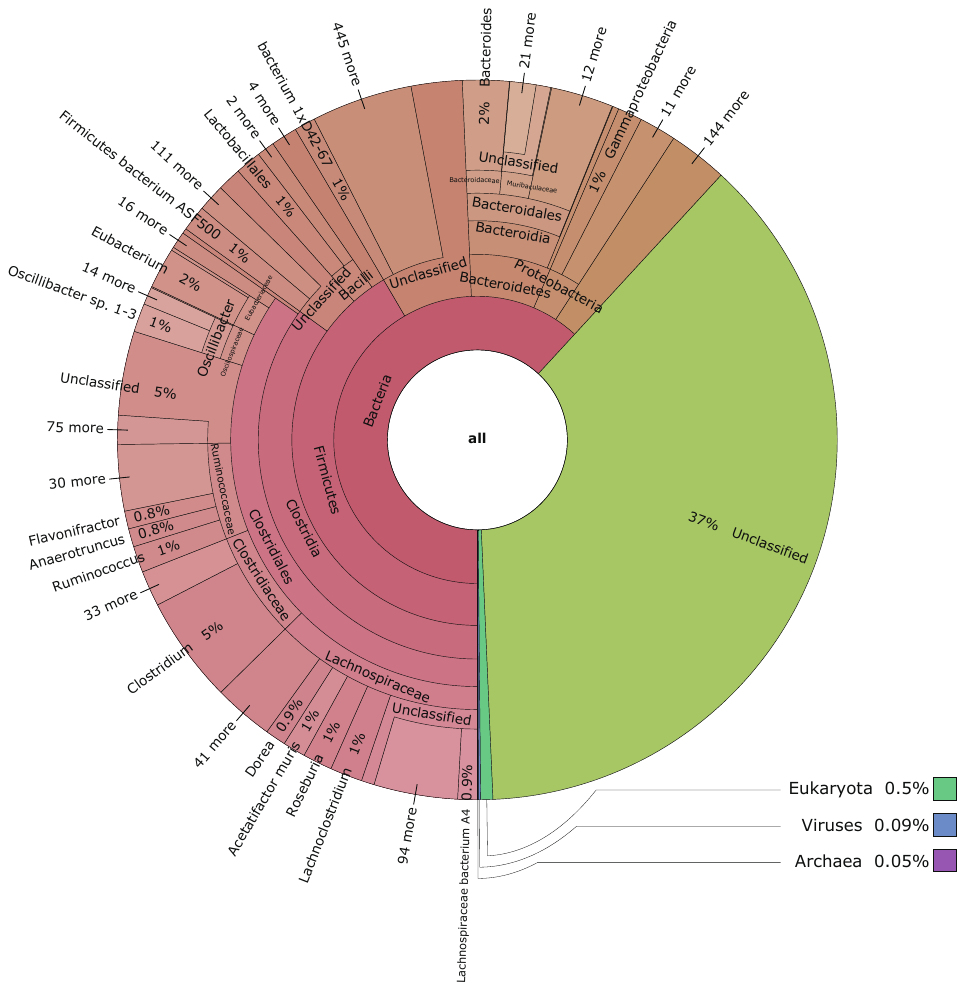

**Supplementary Figure 3** A General View of the Taxonomic Composition of EMGC in a Krona Pie Chart.

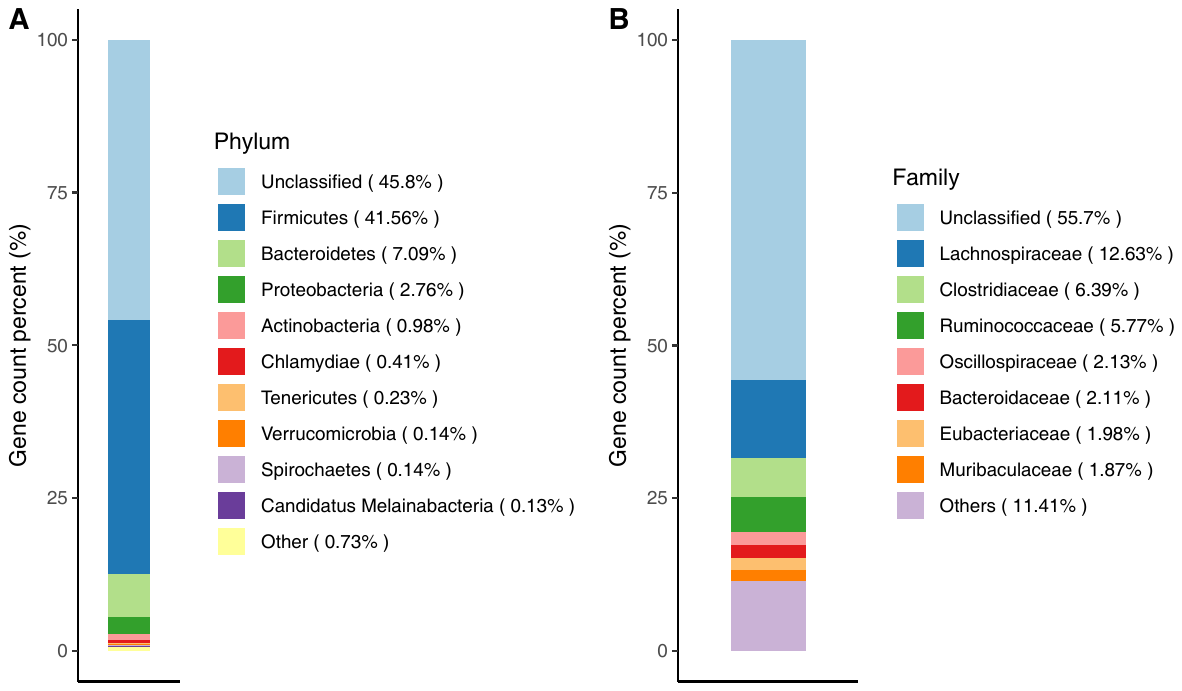

**Supplementary Figure 4** Stacked Bar Plots showing the Distribution of the Taxonomic Annotation of the EMGC at the Phylum Level (A) and Family Level (B).

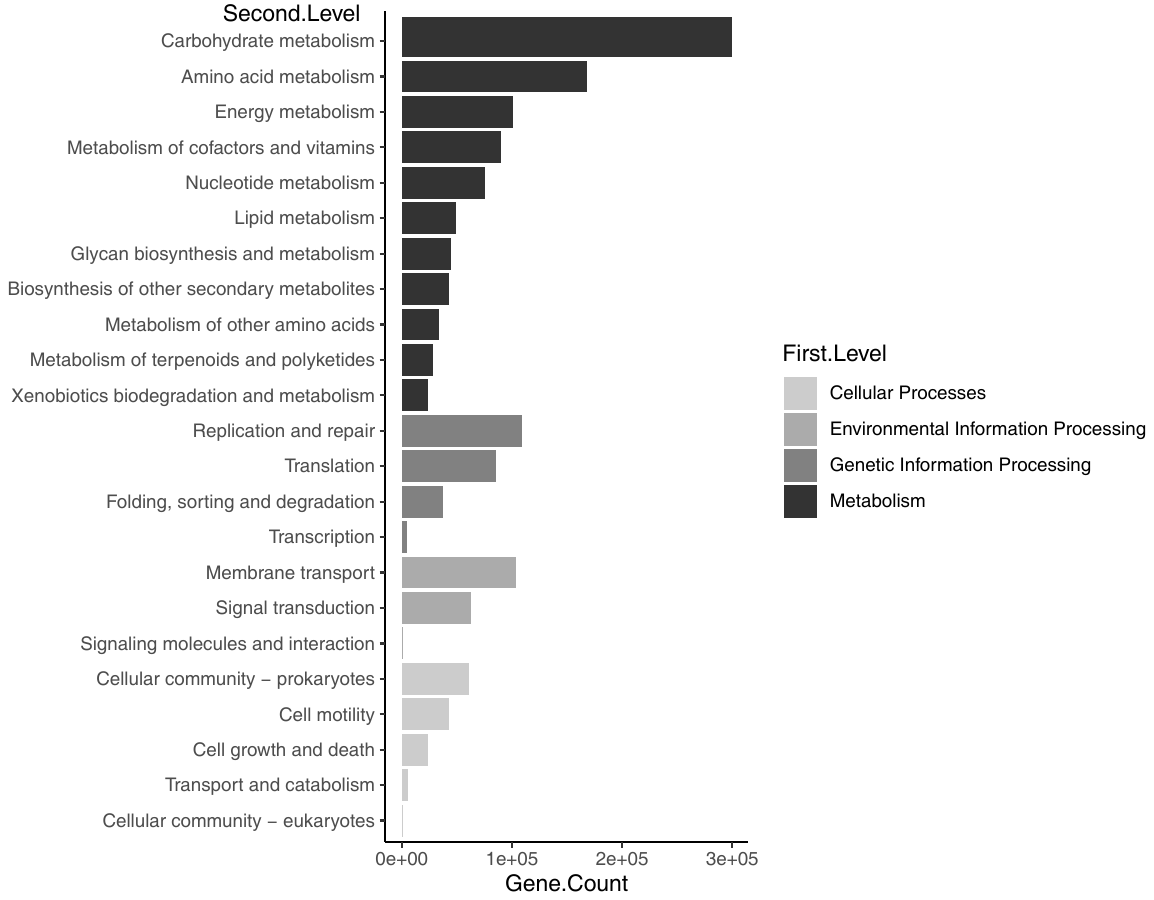

**Supplementary Figure 5** Distribution of annotated functional Pathways according to KEGG in EMGC.

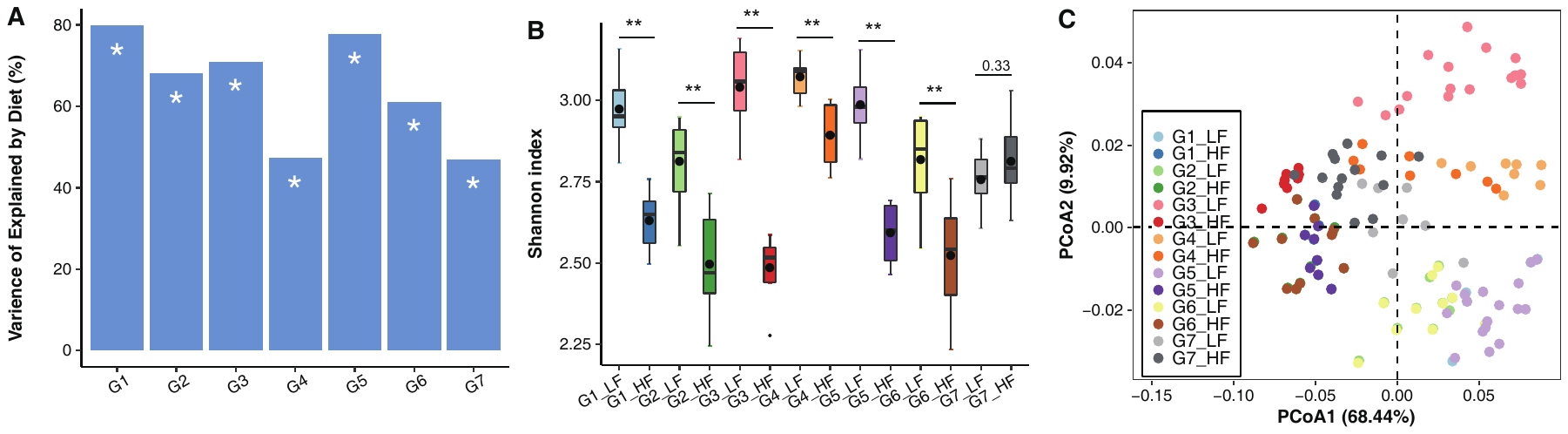

**Supplementary Figure 6** The Influence of Diet on the Composition of Bacterial Functions (KOs) Based on EMGC. (A) PERMANOVA test to estimate the influence of diet on gene functions in the gut metagenomes of different groups. G1: C57/BL6 mice provided by Taconic Denmark (TDK) and hosted in National Institute of Nutrition and Seafood Research of Norway (NIFES); G2: Sv129 mice provided by TDK and hosted in NIFES; G3: C57/BL6 mice provided by the Jackson Laboratory in US (JUS) and hosted by Pfizer-I; G4: C57/BL6 mice provided by Taconic US (TUS) and hosted in Pfizer-I; G5: C57/BL6 mice provided by TDK and hosted in the University of Copenhagen (KU); G6: Sv129 mice provided by TDK and hosted in KU; G7: C57/BL6 mice provided by the Laboratory Animal Center of Sun Yat-Sen University (SYSU). *, p values < 0.001. (B) Boxplot for of the Shannon index of KOs in mice fed both HF and LF diet in each group. **, p values < 0.01, Wilcox rank-sum test. (C) PCoA for the KO profile.

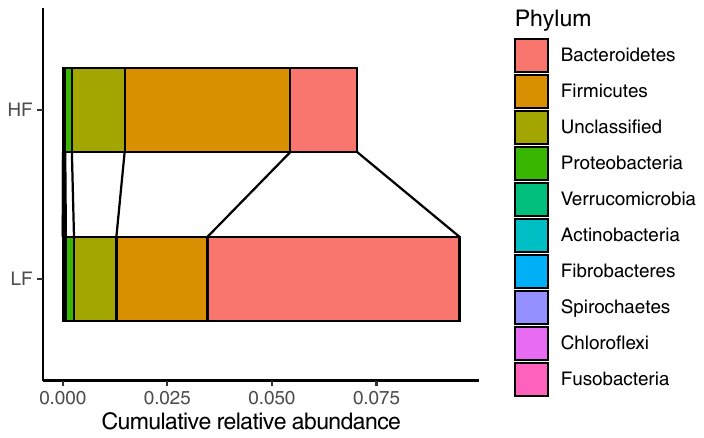

**Supplementary Figure 7** Cumulative relative Abundances of top 10 phyla contributing to Genes related to HF-enriched KOs and LF-enriched KOs.

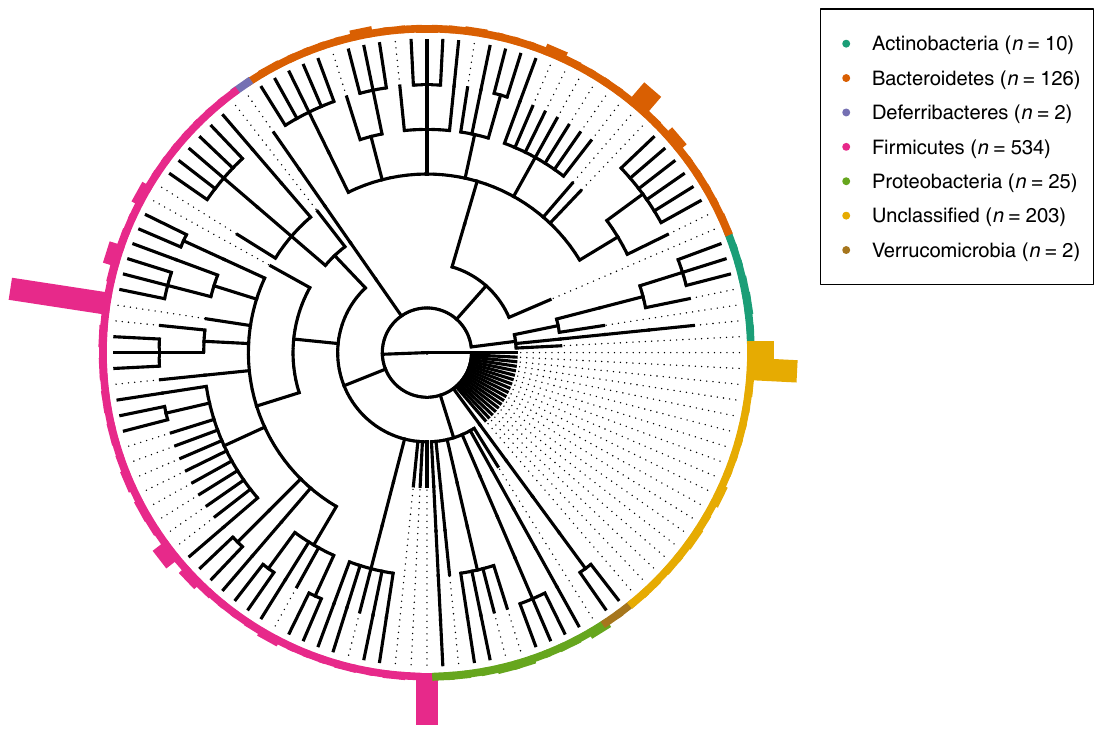

**Supplementary Figure 8** A Cladogram with Bars for the taxonomic Distribution of MGSs clustered from EMGC Gene Profiles.

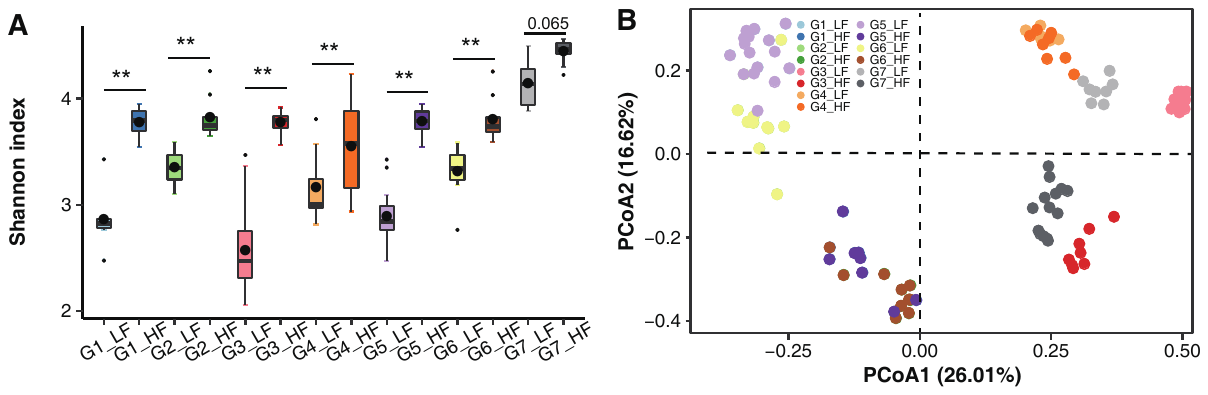

**Supplementary Figure 9** (A) Boxplot showing how different diets in each group affect the Shannon Index based on MGS profiles. **, p value < 0.01, Wilcox rank-sum test (B) PCoA for MGS profile of HF and LF diets in the 7 sample groups.
